# Discovery of a novel coronavirus in Swedish bank voles (*Myodes glareolus*)

**DOI:** 10.1101/2022.02.24.481848

**Authors:** Anishia Wasberg, Jayna Raghwani, Jinlin Li, John H.-O. Pettersson, Johanna F. Lindahl, Åke Lundkvist, Jiaxin Ling

## Abstract

We identified a novel *Betacoronavirus* from bank voles (*Myodes glareolus*) in Grimsö, Sweden. Repeated detection over three years and an overall prevalence of 3.4% suggests the virus commonly occurs in bank voles. Furthermore, phylogenetic analyses indicate the virus belongs to a highly divergent *Embecovirus* lineage predominantly associated with bank voles.

## Text

Bank voles (*Myodes glareolus*) is one of the most common rodent species in Europe and a known reservoir for several zoonotic pathogens, such as *Puumala orthohantavirus* and *Francisella tularensis*. Previous studies have detected various *Alphacoronaviruses* and *Betacoronaviruses* in bank voles in the UK, Poland, Germany, and France (1, 2). Here, following a virome investigation of a Swedish bank voles collected in Grimsö, Sweden, we report two genome sequences of the Grimso virus, along with its evolutionary relationship to other rodent coronaviruses (CoV) and its prevalence over a three-year study period.

Initially, we screened 266 bank voles collected during 2015–2017 at the same site in Grimsö, Sweden (59°43′N, 15°28′ E) for coronaviruses using an in-house PCR method using custom primers targeting the spike gene (Figure 1, and Appendix). We detected nine positive samples, which we further characterized using Sanger sequencing. The detailed method description on bioinformatics and phylogenetic analyses are described in Appendix. We also screened the same samples using a published pan-coronavirus RT-PCR (3), although this failed to detect any coronaviruses.

**Figure 1.**
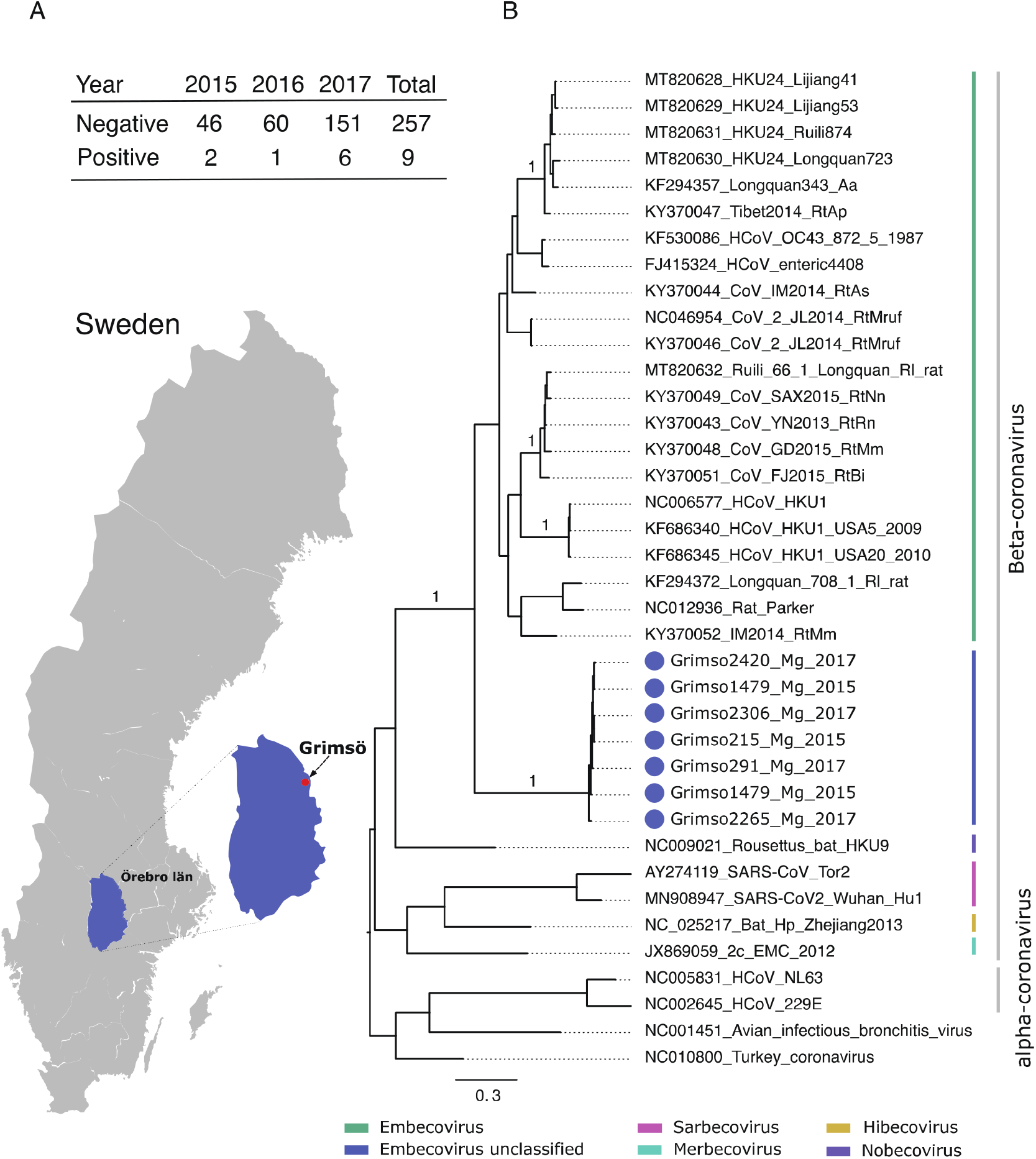
**A)**Geographic map showing the province Örebro, Sweden and the site of sampling (Grimsö) where bank voles were captured. Table demonstrating the prevalence of Grimso virus from 2015 to 2017. **B)** MrBayes midpoint root tree based on the 252 nt of the spike gene. The scale bar indicates the nt substitution per site. The numbers above the branches indicate the posterior probability. Grimso virus samples are highlighted in blue.

Figure 1 shows the location of the study site and the prevalence of the Grimso virus between 2015 to 2017. We obtained partial spike gene sequences (252nt) from seven out of nine positive samples using Sanger sequencing. Pairwise genetic analyses indicated that the partial Grimso virus sequences shared 98.0−100% and 94.0−100% sequence identity at the nucleotide and amino acid levels, respectively. Strikingly, Grimso virus sequences showed less than 60% and less than 50% sequence identity at both the nucleotide and amino acid levels, respectively, with other rodent *Betacoronaviruses*. Bayesian phylogenetic analysis of partial spike gene sequences from 31 reference CoV genome sequences indicates that the Grimso virus sequences form a distinct monophyletic group that cluster with other rodent-borne *Embecoviruses* with strong statistical support (posterior probability = 1.0).

Thereafter, we selected two samples collected in 2015 (Grimso215) and 2017 (Grimso2306) for RNA-sequencing (see Appendix) to characterize the complete genome. We obtained a full-length coronavirus sequence from Grimso215 (31,317 nt) and a near-complete coronavirus sequence from Grimso2306 (98%; 30,767 nt). Figure 2 outlines the whole genome, including seven subgenomic regions, of the Grimso virus, strain Grimso215. The Grimso virus genome encodes for hemagglutinin esterase (HE), spike protein (S), envelop protein (E), membrane protein (M), and nucleocapsid protein (N). We found no significant evidence for recombination in the genome of strain Grimso215. Phylogenetic analyses based on the ORF1b, S and N genes sequences consistently showed that the Grimso215 and Grimso2305 strains fell within known rodent *Betacoronavirus* diversity and formed a distinct lineage within the subgenus *Embecovirus* (Fig 2B).

**Figure 2.**
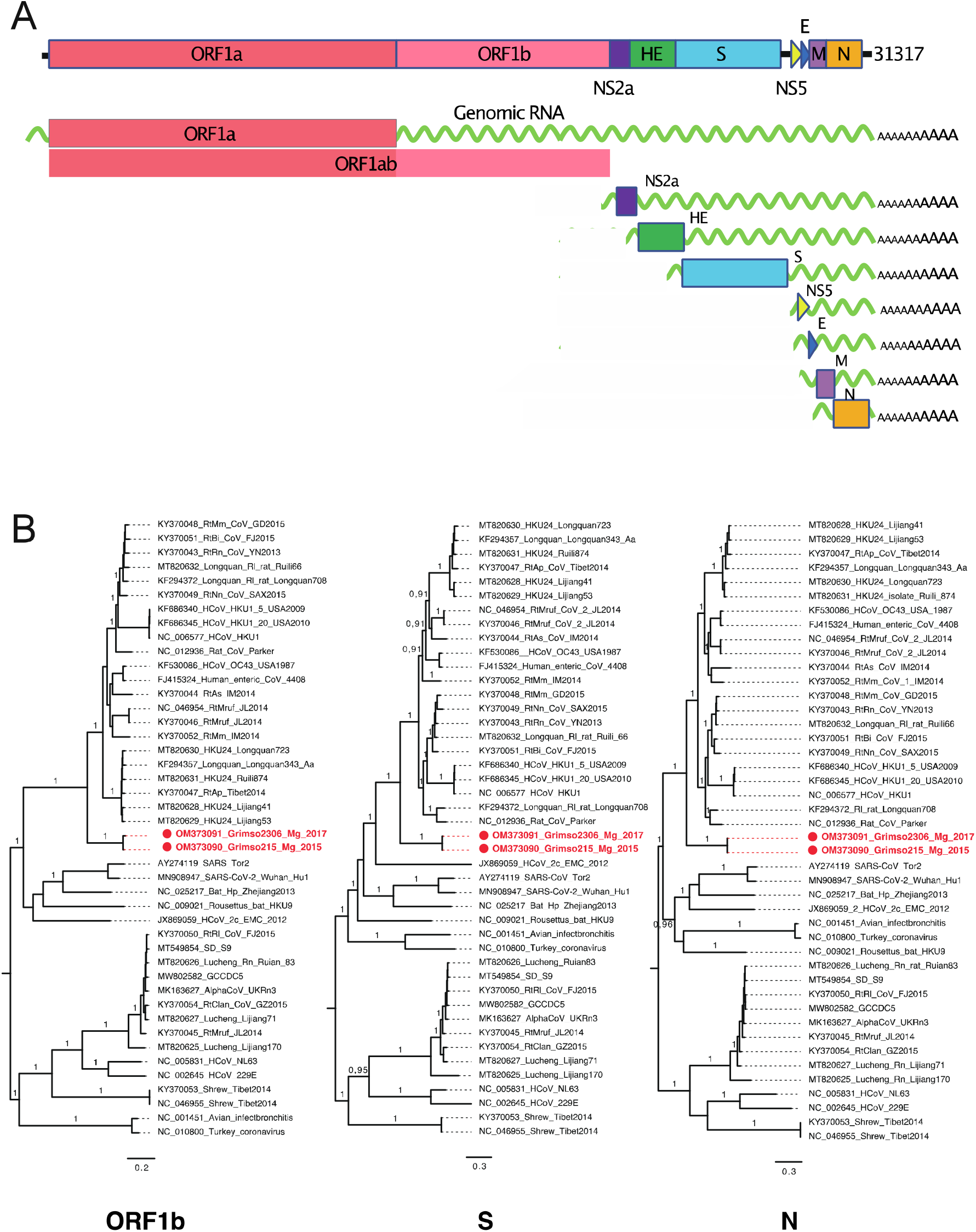
**A)**Genomic RNA and subgenomic mRNAs organizations of Grimso virus, Grimso215 strain. **B)** MrBayes tree based on the complete sequences of ORF1b, S, and N genes of CoVs. The red colour shows Grimso virus. The scale bar indicates the nt substitution per site. The numbers above the branches indicate the posterior probability.

Pairwise sequence identity of Grimso virus (strain Grimso215) with other rodent CoVs ranged from 55−79% at the nucleotide level and 44−67% at the amino acid level, confirming that the Grimso virus is divergent and genetically distinct from earlier described rodent CoVs (Table 1). The genomes of Grimso215 and Grimso2306 strains shared 95.6% sequence identity at the nucleotide level, with 1,338 site differences. This divergence is notably higher than the expected differences based on a typical substitution rate for coronaviruses of 0.001 substitutions per site per year (4, 5), which under Poisson distribution predicts an accumulation of 61−121 substitutions over three years. This observation suggests that either multiple strains of Grimso-like viruses are co-circulating in bank voles in Grimsö or that these viruses are transmitted regularly to bank voles from other species. Nevertheless, with a prevalence of around 3.4% (9/266), we hypothesize that Swedish bank voles are competent hosts for the Grimso virus.

**Table 1.**
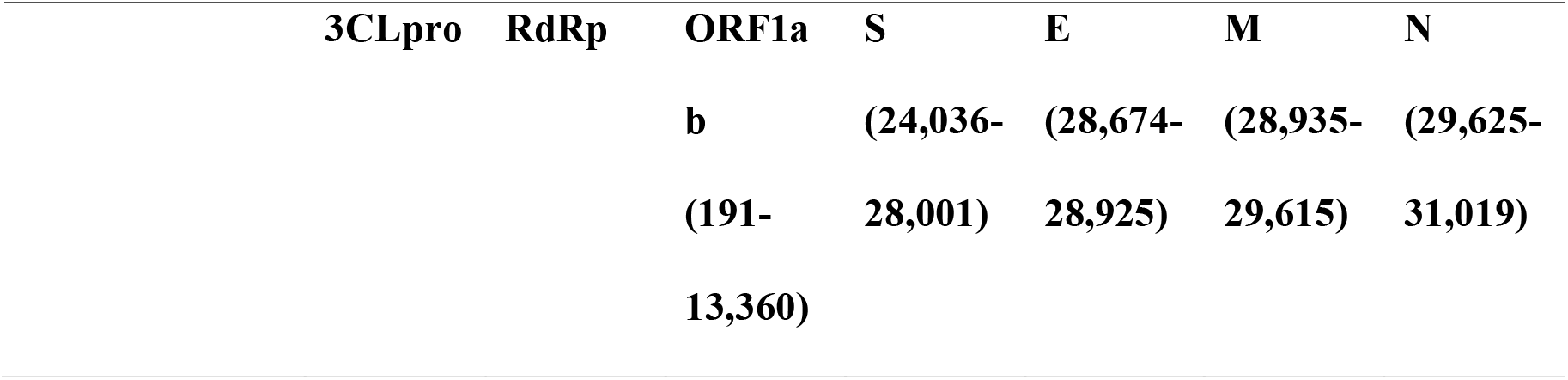
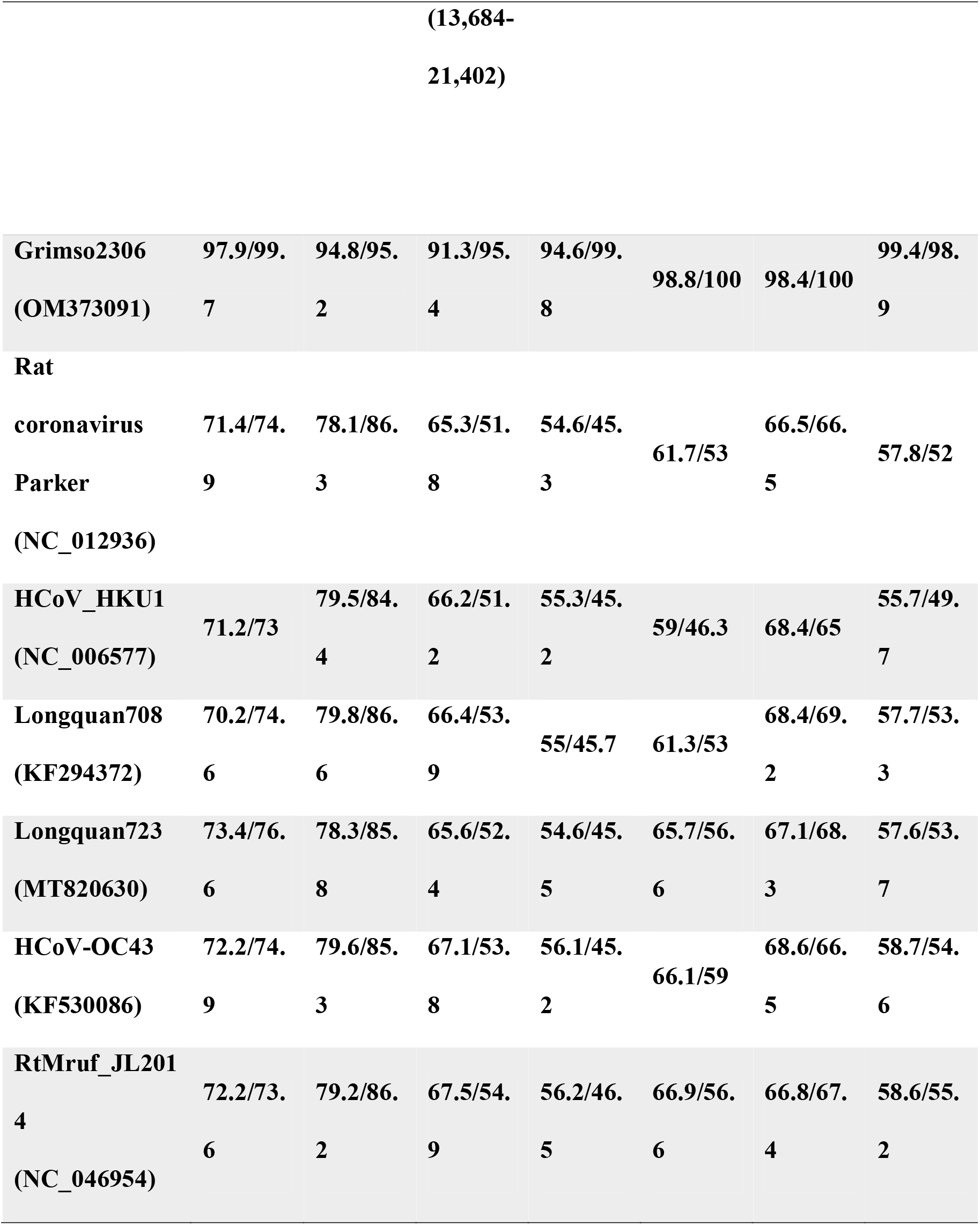
Identity (%) of Grimso215 strain on the genes across the genome (at nucleotide/amino acid levels) as compared to the representatives of known rodent CoVs.

To better understand the evolutionary history of rodent CoVs and divergent Grimso strains, we undertook an additional phylogenetic analysis based on 441 nt of the partial RNA-dependent RNA polymerase (RdRp) gene from 101 CoV genome sequences (Appendix). The results indicated that bank voles harbour at least three virus species across the *Alphacoronavirus* and *Betacoronavirus* genera (Figure S1A and S1B). Specifically, four viruses isolated from bank voles in Germany, Poland, and the UK (6) were closely related to the *Lucheng Rn rat coronavirus* (genus: *Alphacoronavirus*) (Figure S1A). In addition, we also found two distinct *Embecoviruses* associated with bank voles (Figure S1B): 1) a bank vole coronavirus from Germany closely related to *Myodes coronavirus 2JL14* (subgenus: *Embecovirus*) and 2) a cluster of CoVs isolated from bank voles in Germany and France, along with Grimso215 and Grimso2306 strains from this study, which formed separate divergent lineage within the subgenus *Embecovirus* (Figure S1B). Together, these observations suggest a relatively broad geographic distribution of CoVs in bank voles in Europe, indicative of possible long-term host-virus association. Furthermore, as coronaviruses closely related to the Grimso virus have been detected in bank voles elsewhere in Europe, it further supports that this divergent coronavirus infects and circulates in Swedish bank voles.

## Conclusion

We identified a highly divergent *Embecovirus*, named the Grimso virus, in Swedish bank voles. Our analyses suggest that multiple distinct viral strains co-circulate in this population, although further investigation will be necessary to fully understand the transmission ecology. Furthermore, we found that closely related coronaviruses are broadly distributed across Europe and exclusively associated with bank voles and other vole species, indicating that bank voles are likely natural reservoirs of the Grimso virus. While the potential threat posed by the virus to human and animal health is unknown, our findings underscore the importance of longitudinal surveillance of CoVs in wild rodents in advancing current knowledge on the ecology of CoVs in reservoir populations.

## Acknowledgment

We thank Petter Kjellander, Ulrika Alm Bergvall, Per-Erik Lindgren and Jan Chirico for access to the rodent sample collection. We thank Anas Alomar and Magdalena de Arriba Sanchez de la Campa for laboratory assistance. This study was supported by Börjesson E o R stipends awarded to JL. JHOP is funded by the Swedish research council FORMAS (grant no: 2015-710) and VR (grant no: 2020-02593) and JFL by the Swedish research council FORMAS (grant no: 2016-00364). We thank funding resources from European Union’s Horizon 2020 research innovation program under grant no. 874735 (VEO), from The Swedish Research Council (VR: 2017-05807) and by SciLifeLab, Pandemic Laboratory Preparedness (LPP1-007).

## References

1. Tsoleridis T, Onianwa O, Horncastle E, Dayman E, Zhu M, Danjittrong T, et al. Discovery of Novel Alphacoronaviruses in European Rodents and Shrews. Viruses. 2016 Mar 18;8(3):84.

2. Monchatre-Leroy E, Boue F, Boucher JM, Renault C, Moutou F, Ar Gouilh M, et al. Identification of Alpha and Beta Coronavirus in Wildlife Species in France: Bats, Rodents, Rabbits, and Hedgehogs. Viruses. 2017 Nov 29;9(12).

3. Tong S, Conrardy C, Ruone S, Kuzmin IV, Guo X, Tao Y, et al. Detection of novel SARS-like and other coronaviruses in bats from Kenya. Emerg Infect Dis. 2009 Mar;15(3):482–5.

4. Forni D, Cagliani R, Pontremoli C, Clerici M, Sironi M. The substitution spectra of coronavirus genomes. Brief Bioinform. 2022 Jan 17;23(1).

5. Jackwood MW, Hall D, Handel A. Molecular evolution and emergence of avian gammacoronaviruses. Infect Genet Evol. 2012 Aug;12(6):1305–11.

6. Tsoleridis T, Chappell JG, Onianwa O, Marston DA, Fooks AR, Monchatre-Leroy E, et al. Shared Common Ancestry of Rodent Alphacoronaviruses Sampled Globally. Viruses. 2019 Jan 30;11(2).

