## appendix for "Discovery of a novel coronavirus in Swedish bank voles (*Myodes glareolus*)"

**The study**

**Technical Appendix**

**Additional methods and Details**

**Rodent samples collection**

A total of 450 bank voles were sampled at the same site in Grimsö, Sweden (59°43′N, 15°28′E) between 2015 to 2017. All trapping and sampling were approved by Animal Experiment Ethical Committee, Umeå (Reference: A12-14), and followed the Swedish Board of Agriculture regulations. Animal species were identified in the laboratory and lung tissues were harvested and subsequently stored at −80°C until further investigation, including molecular species identification as described in (1).

**RNA extraction and Reverse Transcription-PCR (RT-PCR)**

Total RNA was extracted using Qiagen RNeasy mini kit (Qiagen, Hilden, Germany). Specific primers targeting the spike protein gene (CoVF: 5’-Ggtcaaactactgaatttattg-3’, CoVR: 5’-Aatccatcagaaccaacgac-3’) were designed and used for screening coronaviruses in 266 bank voles captured from Grimsö between 2015 to 2017. Positive samples were sent for Sanger sequencing at Macrogen Europe (https://dna.macrogen-europe.com/eng/).

**Next-Generation Sequencing and sequence assembly**

Two positive RNA samples extracted from lung tissues (Grimso215 and Grimso2306) were sent for RNA-seq, at the Illumina NovaSeq 6000 sequencing platform, from Novogene Hong Kong (<https://en.novogene.com/)>. The number of raw reads reached around 50 million pair-end reads of 150 base-pairs (bp). A data analysis pipeline was used to trim and assemble the reads from the sequencing results as described in (3). We obtained 86,322,748 (96.89% of raw reads) and 105,068,356 (98.81% of raw reads) clean pair-end reads of 150 base-pairs (bp) after filtering from Grimso215 and Grimso2306 samples, respectively. Full-genome and subgenomic sequences of Grimso virus Grimso215 strain were obtained through *de novo* assembly using Trinity v2.13.2 with default settings (4). Using de-novo assembled full genome sequence as a reference, we mapped the reads from the two samples separately (Grimso215: 104,891 reads; Grimso2306: 2700 reads). As a result, we obtained a complete and a near-complete coronavirus genome sequences from Grimso215 strain (100%; 31,317 nt, with a mean coverage value of 502) and Grimso2306 strain (98.2%; 30,767 nt, with a mean coverage of 12.9), respectively. The genome sequences are available via NCBI GenBank (accession numbers: OM373090 and OM373091).

**Phylogenetic analysis**

For the recombination and phylogenetic analyses, the reference rodent CoVs were downloaded from the NCBI RefSeq database, and multiple sequence alignment was obtained using MAFFT v7.490, which was refined using trimAl v.1.4.1 (5, 6). Pairwise genetic distances were obtained using Geneious Prime v.2019.2.1. Potential recombination events were detected by using Simplot v. 3.5.1 and RDP3 (7). We assembled six multiple nucleotide sequences alignments for the tree inference, including A) 31 partial spike protein genes sequences from NCBI RefSeq viral database and 7 from this study (Figure 1), B) 30 partial RdRp genes sequences from alphacoronavirus genome sequences retrieved from the database (Figure S1A); C) 69 partial RdRp gene sequences from betacoronavirus genome sequences retrieved from the database and 2 sequences from this study (Figure S1.B); D) ORF1b gene, E) spike protein gene, and F) nucleocapsid protein gene from 42 coronavirus genome sequences from the database and 2 genome sequences from this study (Figure 2. B). The substitution model was determined by using jModelTest2 (8). Phylogenetic trees were built using MrBayes rooted on the midpoint (9). All computational calculations from this study were done using the UPPMAX service from Uppsala University (<https://www.uppmax.uu.se/)>.

**Protein domain analysis**

We also examined the spike protein, hemagglutinin esterase and possible receptor usages by searching in the blastx. It had only specific hits in the S2 region but not in the S1 or receptor-binding regions (Figure S2). Due to the high divergence and the lack of isolated live virus, we could not yet identify the host receptor usage for this novel rodent CoV. However, we cannot neglect that the Grimso215 virus might pose a zoonotic threat to livestock or humans.

References

1.	Ling J, Sironen T, Voutilainen L, Hepojoki S, Niemimaa J, Isoviita VM, et al. Hantaviruses in Finnish soricomorphs: evidence for two distinct hantaviruses carried by Sorex araneus suggesting ancient host-switch. Infect Genet Evol. 2014 Oct;27:51-61.

2.	Tong S, Conrardy C, Ruone S, Kuzmin IV, Guo X, Tao Y, et al. Detection of novel SARS-like and other coronaviruses in bats from Kenya. Emerg Infect Dis. 2009 Mar;15(3):482-5.

3.	Ling J, Persson Vinnersten T, Hesson JC, Bohlin J, Roligheten E, Holmes EC, et al. Identification of hepatitis C virus in the common bed bug - a potential, but uncommon route for HCV infection? Emerg Microbes Infect. 2020 Dec;9(1):1429-31.

4.	Grabherr MG, Haas BJ, Yassour M, Levin JZ, Thompson DA, Amit I, et al. Full-length transcriptome assembly from RNA-Seq data without a reference genome. Nat Biotechnol. 2011 May 15;29(7):644-52.

5.	Capella-Gutierrez S, Silla-Martinez JM, Gabaldon T. trimAl: a tool for automated alignment trimming in large-scale phylogenetic analyses. Bioinformatics. 2009 Aug 1;25(15):1972-3.

6.	Katoh K, Standley DM. MAFFT multiple sequence alignment software version 7: improvements in performance and usability. Mol Biol Evol. 2013 Apr;30(4):772-80.

7.	Martin DP, Lemey P, Lott M, Moulton V, Posada D, Lefeuvre P. RDP3: a flexible and fast computer program for analyzing recombination. Bioinformatics. 2010 Oct 1;26(19):2462-3.

8.	Darriba D, Taboada GL, Doallo R, Posada D. jModelTest 2: more models, new heuristics and parallel computing. Nat Methods. 2012 Jul 30;9(8):772.

9.	Ronquist F, Huelsenbeck JP. MrBayes 3: Bayesian phylogenetic inference under mixed models. Bioinformatics. 2003 Aug 12;19(12):1572-4.

**Figure S1** MrBayes tree based on 441nt of partial RdRp gene of CoVs. **A)** phylogeny of alphacoronaviruses and **B)** betacoronaviruses. The red colour shows CoVs carried by bank voles across Europe. The scale bar indicates the nt substitution per site. The numbers above the branches indicate the posterior probability.

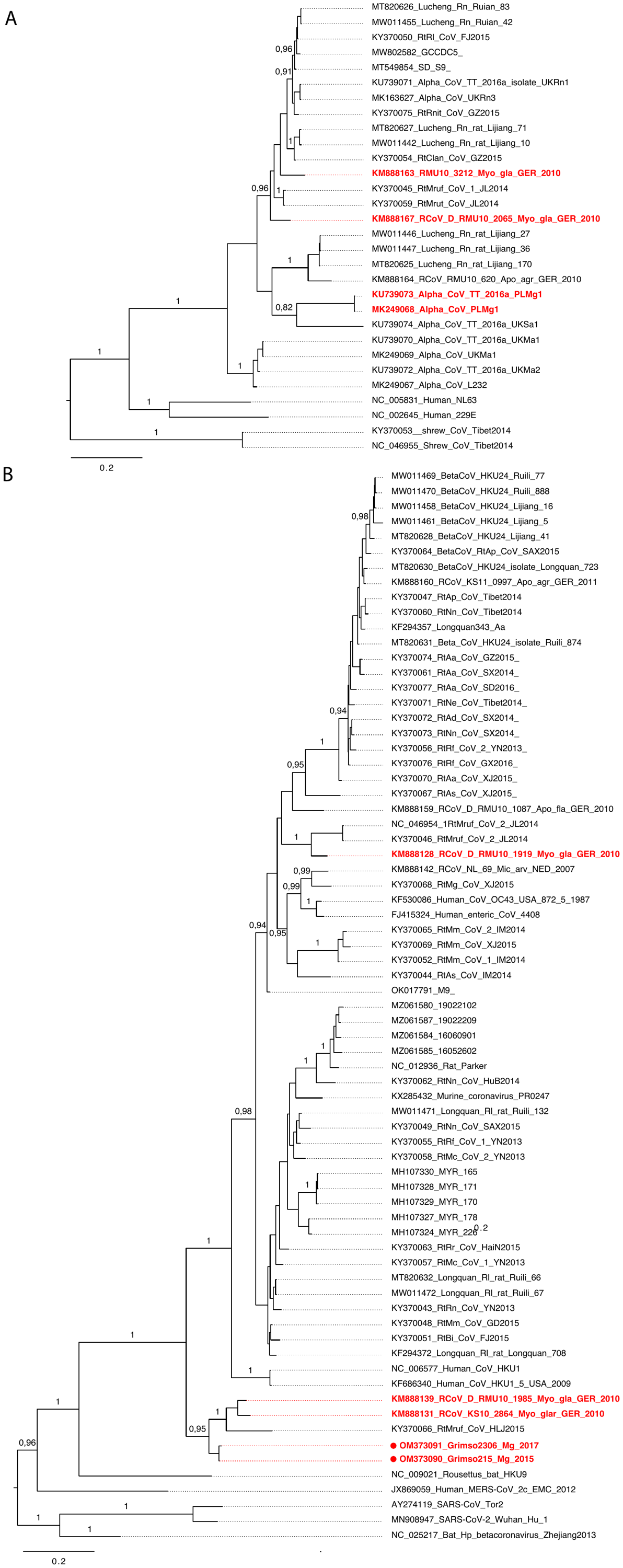

**Figure S2** Conserved domains on the hemagglutinin esterase (A) and spike (B) of Grimso virus.

A

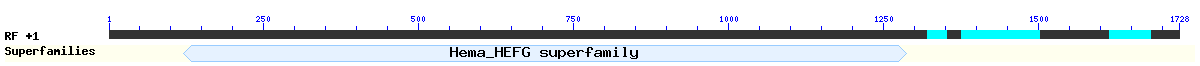

B

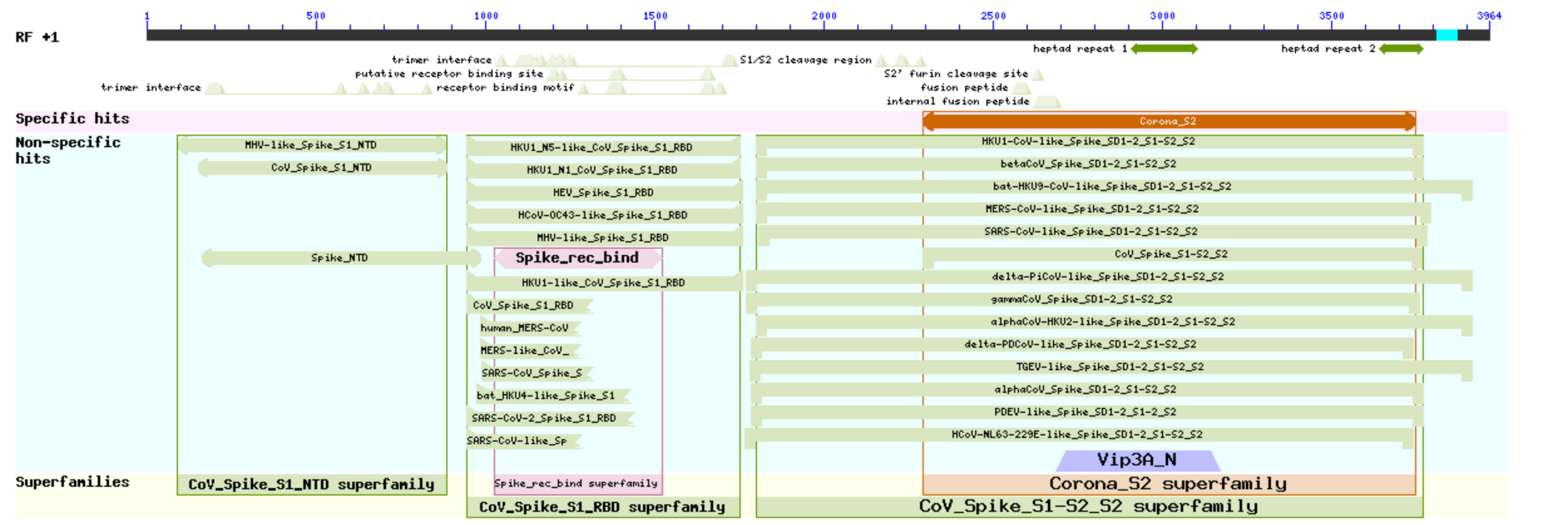
